# Detection of pharmaceutical contamination in amphipods of Lake Baikal by the HPLC-MS method

**DOI:** 10.1101/2024.06.26.600442

**Authors:** Tamara Yu. Telnova, Maria M. Morgunova, Sophie S. Shashkina, Anfisa A. Vlasova, Maria E. Dmitrieva, Victoria N. Shelkovnikova, Ekaterina V. Malygina, Natalia A. Imidoeva, Alexander Yu. Belyshenko, Alexander S. Konovalov, Evgenia A. Misharina, Denis V. Axenov-Gribanov

## Abstract

Pollution by active ingredients is one of the most significant and widespread forms of pollution on Earth. Medicines can have a negative impact on ecosystems, and contamination can have unpredictable consequences. An urgent and unexplored task is to study the Lake Baikal ecosystem and its organisms for the presence of trace concentrations of active pharmaceutical ingredients. Our study aimed to conduct a qualitative analysis of active pharmaceutical ingredients, and quantitative analysis of ibuprofen in endemic amphipods of Lake Baikal, using methods of high-performance liquid chromatography and mass spectrometry (HPLC-MS). Acetylsalicylic acid (aspirin), ibuprofen, acetaminophen, azithromycin, dimetridazole, metronidazole, amikacin, spiramycin, and some tetracycline antibiotics were detected in the studied littoral amphipods. We also detected different annual loads of active pharmaceutical ingredients on amphipods. Using the multiple reaction monitoring (MRM) mode mentioned in GOST International Technical Standards, we detected molecules, fragmented as amikacin, chlortetracycline, doxycycline, oxytetracycline, dimetridazole, metronidazole and spiramycin. Thus, we first revealed that invertebrates of Lake Baikal can encounter pharmaceutical contaminants in the environment.

## INTRODUCTION

Pollution of water bodies by various toxicants, pesticides, or drugs is one of the most significant environmental problems (Rathi, 2021; Khan et al, 2022). It is known that drugs can have negative effects on ecosystems, and pollution can cause unpredictable consequences in both the short and long term (Khan et al, 2022).

Pollution from active ingredients is closely linked to the increasing consumption of drugs by people (Vaudin et al., 2022). This is fueled by such factors as demographic aging, high prevalence of chronic diseases, and the availability and emergence of new drugs. The nonregulated consumption of drugs by mankind leads to penetration of drugs into domestic wastewater in unchanged form or as active metabolites (Grela et al., 2021).

Antimicrobial, antitumor, and antiviral drugs are top sellers in the global pharmaceutical market. Antibiotics are recognized as a group of drugs acting as environmental pollutants (Bilal et al., 2020). Previous studies showed the presence of antibiotics in rivers (Johnson et al., 2015; Zhang et al., 2020), lakes (Yang et al., 2018; Tran et al., 2019), groundwater (Wu et al., 2022), invertebrates (Liu et al., 2018), and other environments. The mass and uncontrolled use of antibiotics contributes to a decrease in their effectiveness and the development of antibiotic resistance (Martinez, 2009; Kotwani et al., 2021), increasing the burden on natural ecosystems.

Several studies showed that drugs such as neomycin, trimethoprim, sulfamethoxazole and enrofloxacin have a high degree of toxicity against test cultures of *Vibrio fischeri, Daphnia magna, Moina macrocopa* and *Oryzias latipes*. Sulfamethazine, oxytetracycline, chlortetracycline, sulfadimethoxine and sulfathiazole are moderately toxic to organisms. Ampicillin and amoxicillin are the least toxic to the above organisms (Park and Choi, 2008).

Drugs related to the non-steroidal anti-inflammatory class are also known to be toxic to inhabitants of aquatic ecosystems (Rastogi et al., 2021; da Silva et al., 2022). Diclofenac, ibuprofen, acetylsalicylic acid, naproxen, and acetaminophen have been reported to be present in different water bodies (Aydin et al., 2019; Castro-Pastrana et al., 2020) as pharmaceutical pollutants. For example, acetaminophen has been detected in wastewater in Europe (Ternes, 1998), USA (Kolpin et al., 2002), and the River Tyne (UK) (Gomez-Olivan et al., 2014). Ibuprofen has been detected in the Llobregat River (Spain) (Ginebreda et al., 2010) as well as in the Pearl River Delta (South China) (Zhao et al., 2010) and in other publications.

In this focus, the effects of pharmaceutical pollution on ancient aquatic ecosystems can be dangerous, little studied and unpredictable. As hypothesized, active ingredients of drugs can reduce the biological diversity of endemic organisms that are highly sensitive to pharmaceutical agents (Néstor and Mariana, 2019). Lake Baikal is one of ancient water bodies. It is a lake of tectonic origin in southeastern Siberia. The lake age is typically cited in the literature as 25-30 million years. However, extensive geological studies suggest that the geological age of Baikal is not less than 60 million years (Matz and Efimova, 2017). Baikal is the deepest lake on the planet and the largest natural reservoir of freshwater (Rusinek et al, 2012). Currently, Lake Baikal is home to more than 2 600 animal species with high rate of endemism (Timoshkin et al., 2016). The Baikal ecosystem is characterized by its structural and ecological complexity.

An urgent and unexplored task is to study the Lake Baikal ecosystem and its organisms for the presence of trace concentrations of active pharmaceutical ingredients. Notably, no monitoring observations regarding the detection of drugs in Baikal zoobenthos have been conducted to date. Invertebrate animals inhabiting the littoral zone of the lake are the first to encounter different pollutants and can accumulate them, thereby facilitating the migration of active ingredients along the food chains.

The aim of this study was to assess the contamination of Lake Baikal amphipods with active pharmaceutical ingredients using high-performance liquid chromatography and mass spectrometry.

## MATERIALS AND METHODS

Amphipods (Amphipoda, Crustacea) were chosen as the subject of our study. Amphipods are one of the most diverse groups of organisms in Lake Baikal, with taxonomic diversity and different ecological preferences. To date, the endemic fauna of Baikal amphipods is represented by more than 350 endemic species and subspecies (Rusinek et al, 2012).

For this study, we selected amphipods of the species *Eulimnogammarus verrucosus* (Gerstf., 1858). Samples were collected with a hydrobiological net on the stone littoral at 0.1-0.5 m depth near Bolshoye Goloustnoye settlement in August 2020 and 2022. Amphipods were fixed at the sampling site into disposable Eppendorf microtubes and frozen at -20°C.

In the laboratory, each individual was weighed using analytical scale (Sartogosm CE224-C, Russia). The wet weight of the amphipods was 0.1-0.7 g. The average weight of amphipods was 0.28 g. The samples were further homogenized using vibration ball mill in the presence of acetonitrile at the ratio of 1:10 (1 – weight of amphipods, g; 10 – volume of acetonitrile, mL). Each sample contained one individual crustacean. The obtained homogenate was centrifuged for 10 min at 3 000 rpm (Armed LC04B Russia). Then 800 μl of supernatant was transferred to microtubes and protein precipitation was performed by adding 10% trichloroacetic acid solution in a volume of 80 μl. Then microtubes were centrifuged again for 10 minutes at 16 000 rpm (Biosan Microspin-12, Latvia). The prepared samples were stored at +4°C. Prior to analysis, the samples were filtered through a 13 mm diameter, 45 μm pore size syringe filter with PVDF membrane, and then 100 μl of the filtrate was transferred into chromatographic vials. The extracts were diluted with 900 μl of acetonitrile.

In this study, we prepared standardized samples of active pharmaceutical ingredients on our own by extracting them from drugs. For qualitative analysis, solutions of aspirin, acetaminophen, ibuprofen, tetracycline, and azithromycin were made from crushed tablets. Analytical standards of ibuprofen (11559-2020, NCAS, Russia) and azithromycin (11570-2020, NCAS, Russia) were used as additional reference positive control. The reference positive control of ibuprofen was used in concentration of 72 ng/mL, and added to extracts of amphipods. Methyl alcohol served as the solvent. The solutions prepared from analytical standards and from crushed tablets were used to determine correct ionization and fragmentation parameters for molecules of the analyzed active ingredients. Additionally, these solutions were used to assess the suitability of chromatographic systems and to establish retention times. To avoid false-positive results, we used the method involving the analysis of natural sample, analytical control and the analysis of a sample with a standard control added to it. The negative control was a mixture of solvents and rinses used to rinse Eppendorf tubes and filters.

A qualitative analysis of the content of other active ingredients was conducted using the multiple reaction monitoring (MRM) mode. The MRM transitions of the studied drugs were described using GOST standards (Gosudarstvennyy Standart, a set of international technical standards maintained by the Euro-Asian Council for Standardization, Metrology and Certification for the Commonwealth of Independent States) employed for control of active ingredients in composition of food products and food raw materials.

Table 1 shows the precursor and product ions, and relevant MRM transitions of all analyzed components, GOST standards are also listed.

**Table 1.** MRM transitions of active ingredients of drugs.

| № | Active pharmaceutical ingredient | Precursor ions (m/z) | Product ions (m/z) | Reference |
| --- | --- | --- | --- | --- |
| 1 | Acetaminophen | 152.0 | 110.0 | Martignac et al., 2013 |
| 2 | Amikacin | 586.3 | 425.2 | GOST 32798-2014 |
| 3 | Amoxicillin | 366.1 | 114.1; 208 | GOST 34533-2019 |
| 4 | Ampicillin | 350.1 | 192.1; 106.1 | GOST 34533-2019 |
| 5 | Apramycin | 540.3 | 217.1 | GOST 32798-2014 |
| 6 | Aspirin | 179.0 | 136.8 | Polagani et al., 2012 |
| 7 | Azithromycin | 749.6 | 591.6 | Chen L. et al., 2007 |
| 8 | Chlortetracycline | 479.1 | 462.1 | GOST 31694-2012 |
| 9 | Clarithromycin | 748.5 | 590.3; 158.0 | GOST 34136-2017 |
| 10 | Demeclocycline | 465.1 | 430.1 | GOST 31694-2012 |
| 11 | Dimetridazole | 142.1 | 81.1; 96.1 | GOST 34533-2019 |
| 12 | Doxycycline | 445.1 | 410.0 | GOST 31694-2012 |
| 13 | Erythromycin | 734.5 | 158.0; 576.3 | GOST 34136-2017 |
| 14 | Florfenicol | 356.1 | 185.1; 119.1 | GOST 34533-2019 |
| 15 | Gentamicin | 478.3 | 157.1 | GOST 32798-2014 |
| 16 | Hygromycin B | 528.3 | 352.1 | GOST 32798-2014 |
| 17 | Ibuprofen | 205.1 | 161.1 | GOST 32881-2014 |
| 18 | Lincomycin | 407.2 | 359.2; 126.0 | GOST 34136-2017 |
| 19 | Metronidazole | 172.1 | 128.1; 82.1 | GOST 34533-2019 |
| 20 | Oxytetracycline | 461.1 | 444.2 | GOST 31694-2012 |
| 21 | Spiramycin | 422.2 | 100.9; 174.1 | GOST 34136-2017 |
| 22 | Sulfamethazine | 279.1 | 186.1; 124.1 | GOST 34533-2019 |
| 23 | Sulfapyridine | 250.1 | 184.1; 156.1 | GOST 34533-2019 |
| 24 | Tetracycline | 445.0 | 410.0 | GOST 31694-2012 |
| 25 | Thiamphenicol | 353.9 | 290.0; 184.9 | GOST 34533-2019 |
| 26 | Tylosin | 916.5 | 772.1; 174.0 | GOST 34136-2017 |

Screening for the presence of active ingredients in samples of Baikal endemic amphipods was performed using the Agilent Infinity II (2019) chromatography-mass spectrometry system with Agilent 6470B triple quadrupole mass spectrometry detector. Agilent Poroshell C18 chromatographic column 2.1 × 50 mm; mobile phase A – 100% Milli-Q water; mobile phase B – 100% acetonitrile; column temperature – 30°C. Table 2 presents the program for HPLC separation. Mass spectrometer setup program: ion source gas temperature: 300°C; ion source gas flow: 5 l/min; nebuliser: 45 psi; drying gas temperature: 250°C; drying gas flow: 11 l/min; capillary voltage: 3.500 V; sample volume: 1 μl.

**Table 2.** HPLC separation program.

| Time, min | 100 % Milli-Q water, % | 100% Acetonitrile, % | Flow rate, ml/min |
| --- | --- | --- | --- |
| 0.5 | 95 | 5 | 0.4 |
| 1.0 | 85 | 15 | 0.4 |
| 6.0 | 20 | 80 | 0.4 |
| 8.0 | 20 | 80 | 0.4 |
| 9.0 | 95 | 5 | 0.4 |

## STATISTIC ANALYSIS

During the study, over 200 samples of amphipods were collected and analyzed. In this paper we present the first qualitative data characterizing pharmaceutical contamination detected in amphipods species of *E. verrucosus*, collected in the settlement of Bolshoe Goloustnoe in August 2020 (N = 16) and 2022 (N = 44). Each analyzed sample contained one individual crustacean. The analysis of each sample was conducted three times. The statistical processing was performed in the Past software (V4.03) using the ANOVA analysis of variance with the Mann-Whitney criterion. Differences between the mean values of the parameters were considered significant at p ≤ 0.05.

## RESULTS

During the first stage of the study, we estimated the retention times of analytes and achieved simultaneous detection of five analyzed components by selecting chromatographic conditions. The peak retention time for aspirin was 4.6 min, for azithromycin - 2.9 min, for ibuprofen - 5.7 min, for acetaminophen - 0.5 min and for tetracycline - 2.6 min. The variance of retention time was 0.05-0.07 min. Table 2 shows data on the presence of trace concentrations of active pharmaceutical ingredients in amphipods collected near B. Goloustnoye settlement in August 2020. Aspirin was detected in 10 out of 16 analyzed samples. Azithromycin and ibuprofen were found in 9 and 12 samples, respectively.

Then we performed additional screening of drugs whose content in food products is regulated by GOST. The results of these experiments are presented in Table 3. The following precursor and product ions of drugs were detected in all 16 samples of amphipod species *E. verrucosus*: chlortetracycline, doxycycline, oxytetracycline and metronidazole. Selective molecular ions of amikacin, dimetridazole and spiramycin were detected in several samples of amphipods. Molecular ions of dimetridazole were detected in 11 out of 16 samples. Molecular ions of spiramycin were detected in 4 samples. Ions of amikacin was detected in 15 out of 16 samples. Figure 1 shows the chromatograms of metronidazole, including the chromatograms of precursor and product ions.

**Table 3.** The presence of active of pharmaceutical ingredients in samples of amphipods of *E. verrucosus* species collected in August 2020.

| № | Azithromycin | Tetracycline | Acetaminophen | Ibuprofen | Aspirin |
| --- | --- | --- | --- | --- | --- |
| 1 | - | - | - | + | + |
| 2 | - | - | - | - | - |
| 3 | + | + | + | + | + |
| 4 | + | + | + | + | + |
| 5 | - | - | - | - | - |
| 6 | - | + | + | - | - |
| 7 | - | - | - | - | - |
| 8 | + | + | + | - | - |
| 9 | + | + | + | + | + |
| 10 | + | + | + | + | + |
| 11 | + | + | + | + | + |
| 12 | + | + | + | + | + |
| 13 | + | + | + | + | + |
| 14 | + | + | + | + | + |
| 15 | + | + | + | + | - |
| 16 | + | + | + | + | + |
| Total |  |  |  |  |  |
| Uncontaminated samples, % | 31 | 25 | 25 | 31 | 37 |
| Contaminated samples, % | 69 | 75 | 75 | 69 | 63 |

Legend: “+” – means the presence of contaminant

**Table 4.** The presence of active pharmaceutical ingredients, controlled by GOST standards in food products, and detected in samples of amphipods.

| № | Amikacin | Chlortetra<br>cycline | Doxycyclin<br>e | Oxytetracycline | Dimetridazole | Metronidazole | Spiramycin |
| --- | --- | --- | --- | --- | --- | --- | --- |
| 1 | + | + | + | + | + | + | - |
| 2 | - | + | + | + | + | + | - |
| 3 | + | + | + | + | + | + | - |
| 4 | + | + | + | + | + | + | - |
| 5 | + | + | + | + | + | + | - |
| 6 | + | + | + | + | + | + | - |
| 7 | + | + | + | + | + | + | - |
| 8 | + | + | + | + | + | + | - |
| 9 | + | + | + | + | + | + | - |
| 10 | + | + | + | + | + | + | - |
| 11 | + | + | + | + | + | + | - |
| 12 | + | + | + | + | - | + | + |
| 13 | + | + | + | + | - | + | + |
| 14 | + | + | + | + | - | + | - |
| 15 | + | + | + | + | - | + | + |
| 16 | + | + | + | + | - | + | + |
| Total |  |  |  |  |  |  |  |
| Uncontaminated samples, % | 6 | 0 | 0 | 0 | 11 | 0 | 75 |
| Contaminated samples, % | 94 | 100 | 100 | 100 | 69 | 100 | 25 |

Legend: “+” - means the presence of contaminant

**Fig. 1.**
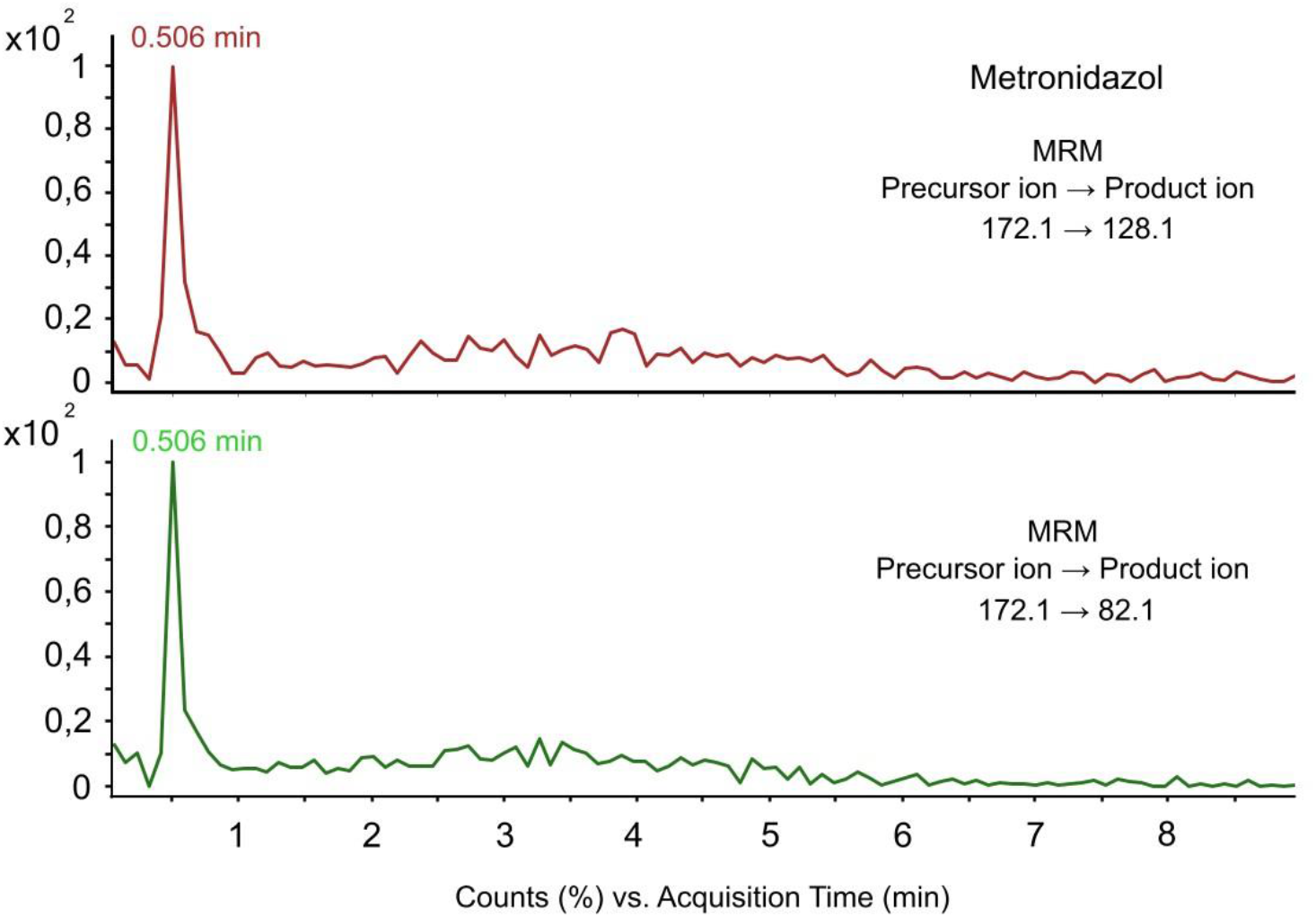
Chromatogram of metronidazole detected in samples of amphipod *E. verrucosus*

At the same time, samples of *E. verrucosus* did not reveal selective molecular ions of the following substances: erythromycin, clarithromycin, tylosin, lincomycin, ampicillin, amoxicillin, florfenicol, thiamphenicol, sulfamethazine, sulfapyridine, gentamicin, hygromycin B, apramycin, and demeclocycline.

The monitoring conducted at the same sampling point in 2020 and 2022, revealed differences in pharmaceutical contamination of amphipod species. Based on the analysis of ibuprofen (Fig. 2), we detected that almost 70% of *E. verrucosus* were contaminated in 2020, and 27% of *E. verrucosus* specimens were contaminated in 2022 (p>0,05). Analysis of other pharmaceutical pollutants using MRM or selective mode was not performed in 2022. The analysis revealed a high correlation between the concentration of ibuprofen and the wet weight of *E. verrucosus* amphipods. The concentration of ibuprofen was found to be higher in small (juvenile) amphipods than in the big (adult) animals (Fig.3). Thus, *E. verrucosus* amphipods with up to 0.2 g wet weight contained ibuprofen in concentrations ranging from 74.71 to 166.38 ng/g. Amphipods with a wet weight of 0.2 to 0.5 g contained ibuprofen in concentrations ranging from 18.50 to 61.75 ng/g. Finally, amphipods with a wet weight above 0.7 g were characterized by ibuprofen concentrations in the range of 14.92-16.98 ng/g.

**Fig.2.**
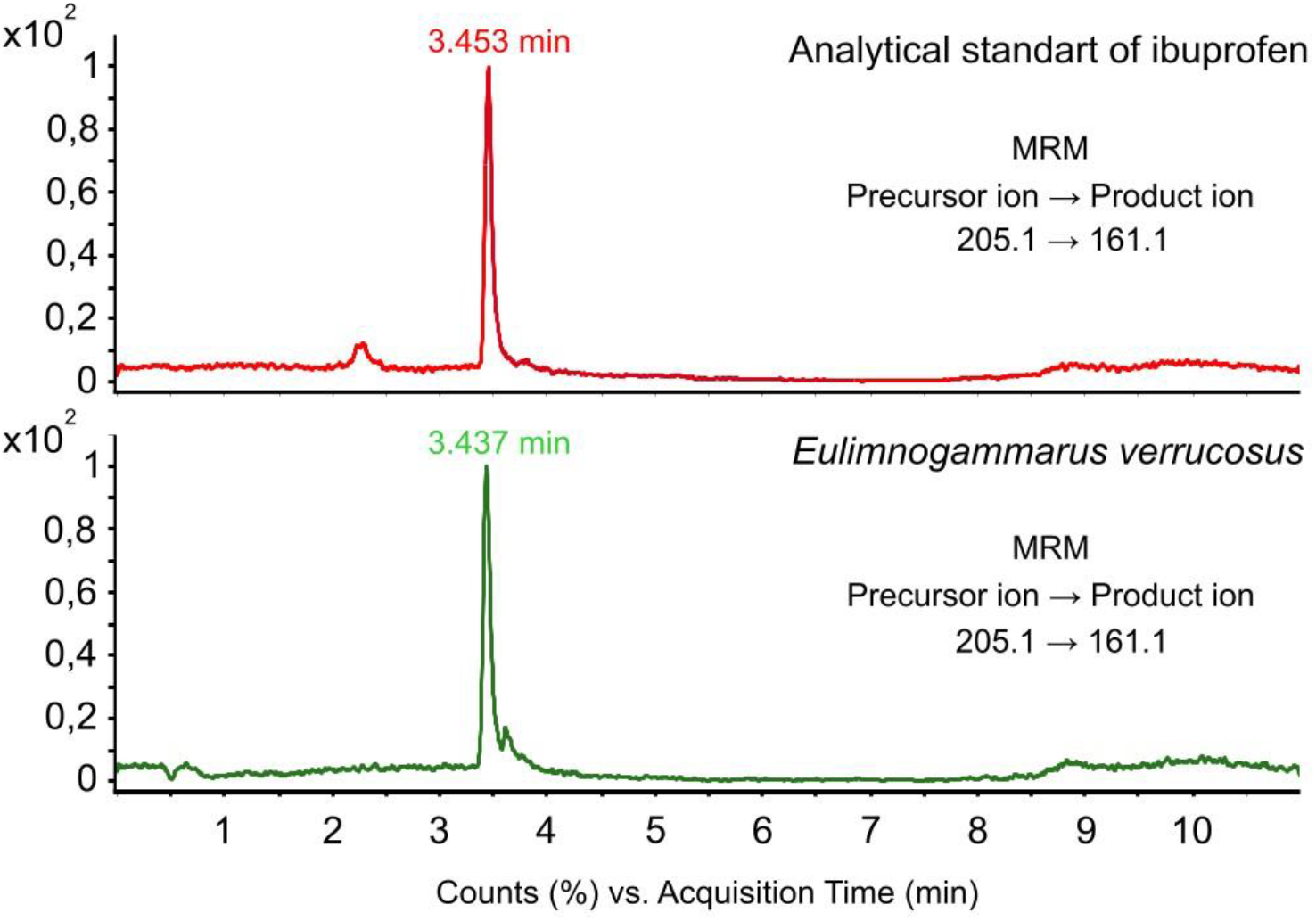
Chromatogram of ibuprofen detected in samples of amphipod *E. verrucosus*

**Fig. 3.**
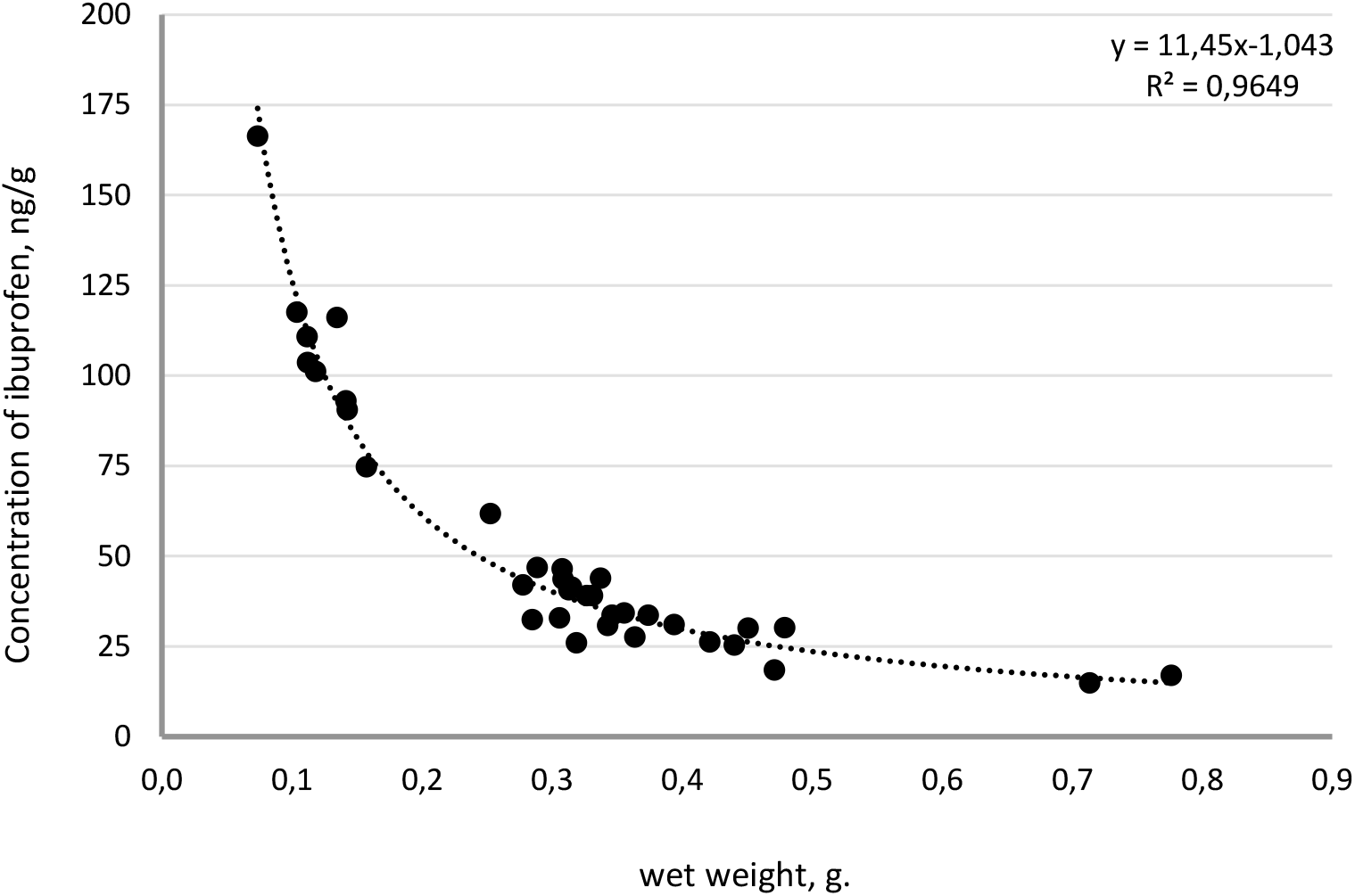
Correlation between concentration of ibuprofen and the wet weight of species *E. verrucosus* amphipods, ng/g.

## DISCUSSION

This study confirmed the hypothesis regarding contamination of Lake Baikal zoobenthos representatives, using amphipods as an example, with active pharmaceutical ingredients, or their selective molecular ions. Contamination of the Lake Baikal ecosystem and its amphipods by active pharmaceutical ingredients is a significant and urgent problem. However, only a few studies have been published to date that characterize pollution of Lake Baikal by active pharmaceutical ingredients.

Studies conducted by Meyer M. F. et al. (2022) found the presence of pharmaceutical agents such as acetaminophen, paraxanthine, caffeine, cotinine, cimetidine, diphenhydramine, phenazone, and sulfachloropyridazine in Lake Baikal water. Other ingredients such as amphetamine, morphine, carbamazepine, tenamphetamine, methamphetamine, thiabendazole, methylenedioxymethamphetamine, sulfamethazine, sulfamethoxazole, and trimethoprim were not detected.

Our study shows that each of 16 samples of *E. verrucosus* collected in B. Goloustnoye settlement in August 2020 contains at least five drug contaminants. Also, ibuprofen was detected in samples collected in 2022 (N=44). Thus, the presence of aspirin, acetaminophen, tetracycline, azithromycin and ibuprofen in amphipods was confirmed. However, different levels of contamination and the absence of quantitative data do not allow us to assume pathways for transformation and migration of active pharmaceutical ingredients through the ecosystem.

The presence of active pharmaceutical ingredients in the endemic amphipods of *E. verrucosus* species can be explained by their entry into the ecosystem through sewage or groundwater. For example, paracetamol (acetaminophen), ibuprofen and azithromycin are of synthetic origin, and they are widely used in medicine, veterinary medicine and households (Freo et al, 2021; Oliver and Hinks, 2021; Bello and Dye, 2023; Chalifoux et al., 2023).

Detection of these molecular ions of drugs in amphipods may indicate their regular entry into domestic wastewater in unchanged form (Mohan et al., 2021). For instance, acetaminophen has been previously detected in water (Meyer et al., 2022), suggesting that this active ingredient is regularly released into the ecosystem of Lake Baikal. The presence of other pharmaceutical contaminants such as dimetridazole, metronidazole, and amikacin may be attributed to successful use of antibiotics in livestock to treat and prevent infectious diseases (Stephens, 1987; Granja, 2013; Motos et al., 2023).

In addition to their synthetic origin, active ingredients of drugs can also be obtained naturally. For example, acetylsalicylic acid is a derivative of salicylic acid, which is the active ingredient in willow (Adamczak, 2019), raspberry, blackberry and blackcurrant extracts (Jeffreys, 2008). Thus, aspirin can enter the water not only with sewage but also with particles of bark and leaves of plants containing salicylate. Thus, amphipods, being phyto- and detritophagous, could consume these natural products with food, as Baikal algae, whose chemistry of natural products is not studied.

Also, some naturally occurring active ingredients can be detected due to symbiotic microorganisms living in the gastrointestinal tract or hemolymph of amphipods. These microorganisms are capable of producing natural compounds. For example, tetracycline is a semi-synthetic antibiotic and can also be produced by actinobacteria related to *Streptomyces aureofaciens* and *S. Rimosus*. These microorganisms are regular symbionts of amphipods (Darken, 1960; Axenov – Gribanov et al., 2016; Quinn, 2020). Notably, oxytetracycline, doxycycline and chlortetracycline identified during the study also belong to the tetracycline group. Spiramycin refers to natural antibiotics in the macrolide group and is synthesized by actinobacteria of species *Streptomyces ambofaciens* (Risdian et al., 2019).

Thus, our study demonstrates that amphipods of Lake Baikal can encounter pharmaceutical contaminants in their environment.

## CONCLUSION

Our study reveals the fact of contamination of Baikal endemic amphipods of *E. verrucosus* species by active pharmaceutical ingredients. The presence of ibuprofen, acetominophen, tetracycline and other pollutants can have a negative impact on endemic organisms of Lake Baikal ecosystem. For this reason, it is necessary to organize comprehensive monitoring studies with the aim of assessing the state of the environment.

## Acknowledgments

*The study was carried out with the financial support of the project of the Ministry of Higher Education and Science of the Russian Federation (project FZZE 2024-0003, FZZE 2024-0011)*.

## Ethical Approval

Not applicable.

## Consent to Participate

Not applicable.

## Consent to Publish

Not applicable.

